# Impaired Myelination in Multiple Sclerosis Organoids: p21 Links Oligodendrocyte Dysfunction to Disease Subtype

**DOI:** 10.1101/2025.01.08.631924

**Authors:** Nicolas Daviaud, Tanmay Mehta, William Holzman, Annie McDermott, Saud A Sadiq

## Abstract

Multiple sclerosis (MS) is an autoimmune inflammatory disease of the central nervous system. The cause of the disease is unknown but both genetic and environmental factors are strongly implicated in its pathogenesis. We derived cerebral and spinal cord organoids from induced pluripotent stem cells (iPSC) from healthy controls as well as from primary progressive MS (PPMS), secondary progressive MS (SPMS) and relapsing-remitting MS (RRMS) patients to investigate and compare oligodendrocyte differentiation and myelination capacity in healthy subjects and MS subtypes. In MS organoids, particularly in PPMS, we observed a decrease in p21 expression associated with a dysregulation of PAK1 and E2F1 expression. In parallel, a decrease in oligodendrocyte maturation was detected in long-term cultured cerebral and spinal cord organoids, especially in PPMS, leading to a reduced myelination capacity. Disruption of astrocyte and neuronal populations was also observed. Our findings demonstrate that in MS, inherent deficits in the p21 pathway may alter glial and neuronal cell populations and may contribute to the disease pathogenesis by reducing the capacity for myelin repair.

**Summary Statement:** Using cerebral and spinal cord organoids derived from multiple sclerosis patients, we found an innate disruption of oligodendrocyte differentiation and myelination capacity as well as excitotoxicity, associated with PAK1 and E2F1-induced p21 dysregulation.

## INTRODUCTION

Multiple sclerosis (MS) is an auto-immune inflammatory disorder that may lead to irreversible neurological disability and cognitive decline (Filippi et al., 2018). It is characterized by widespread focal lesions of primary demyelination in the brain and spinal cord, with variable axonal, neuronal, and astroglial injury. The disease presents primarily as two clinical subtypes: relapsing-remitting MS (RRMS), which accounts for 85-90% of cases which may evolve to secondary progressive MS (SPMS), and primary progressive MS (PPMS) which affects about 10% of cases and is marked by a steady functional decline from disease onset (Lublin and Reingold, 1996; Andersson et al., 1999). The origin, evolution, and physiological basis of MS’s varied phenotypic expressions remain poorly understood in part due to the relative inaccessibility of human brain tissues and limitations of animal models (Ransohoff, 2012). It is now accepted that the development of MS is influenced by genetic factors, with familial relatives of patients, especially first-degree relatives, being more susceptible to developing MS compared to the general population. There is also overwhelming epidemiological evidence that environmental triggers such as infections are important in disease causation (Gouider et al., 2024). This interplay of genetic susceptibility and environmental factors contributes to the multifaceted nature of MS (Willer et al., 2003).

In our previous work, we described cerebral-organoids (c-organoids) as an innovative model to study MS. Using c-organoids from iPSCs of healthy control subjects as well as from PPMS, SPMS and RRMS patients, we showed a decrease of proliferative capacity, notably in progressive forms of MS, associated with a reduction of the progenitor pool and an increase of neurogenesis possibly due to a symmetric shift of the mode cell division. We linked these effects to a strong decrease of p21 expression in PPMS organoids, unrelated to the DNA damage and apoptosis pathway. However, our observations were based on organoids that had been in culture for only 42 days.

We report here the use of organoids cultured for 120 to 150 days in vitro to analyze neural and glial cell differentiation and maturation in MS, as well as the myelination capacity of oligodendrocytes. Using innovative cerebral and spinal cord organoid models, we detected a significant decrease in oligodendrocyte maturation and myelination capacity in MS organoids, especially in its progressive form. Additionally, a defect of astrocyte population and an imbalance of inhibitory/excitatory neurons were found, which are key factors in MS onset and possibly in causing cognitive impairment, physical disability, and fatigue in MS patients. We confirmed that the p21 pathway dysfunction was a critical abnormality in our organoid model of MS, while other Cyclin Dependent Kinase inhibitors (CDKi), such as p16, p27 and p57, expressions did not seem to be affected. Furthermore, p21 regulators E2F1 and PAK1, that participate in oligodendrocyte differentiation and myelination and in regulation of the excitatory/inhibitory balance, were also dysregulated.

In conclusion, this work shows that c-organoids derived from patients with MS can be used as an innovative tool to better understand the genetic basis for phenotypic differences seen in MS.

## MATERIALS AND METHODS

### Patient selection

Peripheral blood mononuclear cell (PBMCs) samples were collected from healthy control subjects and clinically definite MS patients diagnosed according to the revised 2017 McDonald Criteria. All MS patients underwent neurological examination and MRI imaging and were classified as having RRMS, SPMS or PPMS by board-certified neurologists specializing in MS care. All protocols were IRB approved, and all donors provided written informed consent for participation. Human IPS cells were generated from PBMCs from donors blood samples. Other cell lines were obtained from the New York Stem Cell Foundation. Donor information can be found on table 1 and table 2.

**Table 1:** Patients details. Table summarizing patients characteristics such as sex, gender, age, diagnosis, and cell source.

| **Sex** | **Age** | **Diagnosis** | **Source** |
| --- | --- | --- | --- |
| M | 28 | Control | NYSCF |
| M | 23 | Control | Tisch MSRCNY |
| M | 36 | Control | Tisch MSRCNY |
| M | 67 | Control | Tisch MSRCNY |
| F | 52 | Control | Tisch MSRCNY |
| F | 40 | Control | Tisch MSRCNY |
| F | 62 | PPMS | NYSCF |
| M | 61 | PPMS | NYSCF |
| F | 58 | PPMS | Tisch MSRCNY |
| M | 40 | PPMS | Tisch MSRCNY |
| F | 57 | PPMS | Tisch MSRCNY |
| M | 38 | RRMS | Tisch MSRCNY |
| F | 67 | RRMS | Tisch MSRCNY |
| M | 24 | RRMS | NYSCF |
| F | 50 | RRMS | NYSCF |
| M | 60 | SPMS | Tisch MSRCNY |
| M | 48 | SPMS | Tisch MSRCNY |
| F | 54 | SPMS | Tisch MSRCNY |
| F | 34 | SPMS | Tisch MSRCNY |
| F | 67 | SPMS | Tisch MSRCNY |

**Table 2:** Patients distribution. B) Table summarizing the age and gender distribution in our cohort. No significant difference was observed in age (One-way ANOVA, p = 0.3450) or gender distribution among the different conditions (Chi-square, p = 0.7851).

|  | Healthy Controls | PPMS | RRMS | SPMS |  |
| --- | --- | --- | --- | --- | --- |
| Number of patients | 6 | 5 | 4 | 5 |  |
| Age at collection, mean (SD), years | 41 (16) | 55.6 (8.9) | 44.7 (18) | 53 (13) | n.s. |
| Gender distribution | 2 F; 4 M | 3 F; 2 M | 2 F; 2 M | 3 F; 2 M | n.s. |

### Reprogramming CD34^+^ progenitor cells

Human PBMCs were isolated according to the “STEMCELL Integrated Workflow for the Isolation, Expansion, and Reprogramming of CD34^+^ Progenitor Cells”. Briefly, blood samples were collected in heparin-vacutainer tubes from donors in amounts ranging from 8 to 20 ml. CD34^+^ hematopoietic stem and progenitor cells were isolated from peripheral blood using the EasySep RosetteSep Kit (STEMCELL) and expanded in vitro in CD34^+^ expansion media consisting of StemSpan SFEM II and CD34^+^ expansion supplements (STEMCELL). After 7-10 days of culture, 1×10^6^ cells were collected for reprogramming by electroporation using the Epi5 Episomal iPSC Reprogramming Kit (ThermoFisher) and Human CD34^+^ Cell Nucleofector Kit (Lonza) using a Nucleofector 2b Device (Lonza). After electroporation cells were cultured on Cultrex UltiMatrix coated 6-well plates (100 µg/ml, R&D systems) in CD34^+^ expansion media. After 3 days, ReproTeSR (STEMCELL) was added to culture media for 2 more days. On day 7 cells were cultured in ReproTeSR media only. Media was changed daily. After 2-3 weeks, IPS cell colonies were isolated manually and transferred to Cultrex coated 6-well plates containing mTeSR plus media (STEMCELL). Three subclones were created for each IPS cell line.

### Human induced pluripotent stem cells

Human iPSCs were cultured as previously described (Lancaster and Knoblich, 2014; Daviaud et al., 2023). iPSCs were plated in 6-well tissue culture plates coated with diluted Cultrex UltiMatrix (100 µg/ml, R&D Systems) and maintained in mTeSR Plus Culture Media (STEMCELL), supplemented with rock inhibitor Thiazovivin (2 µM, Millipore). Media was then changed daily without rock inhibitor until ready to passage at approximately 70-80% confluence or harvested.

All human pluripotent stem cells were maintained below passage 30 and confirmed negative for mycoplasma using the MycoFluor Mycoplasma Detection Kit (ThermoFisher). iPSCs were regularly evaluated for pluripotency using OCT4, NANOG and SOX2 markers (Figure 1) and were confirmed to be karyotypically normal by G-band testing by a qualified service provider (Cell Line Genetics, Madison, WI, USA) (Figure 1).

**Figure 1:**
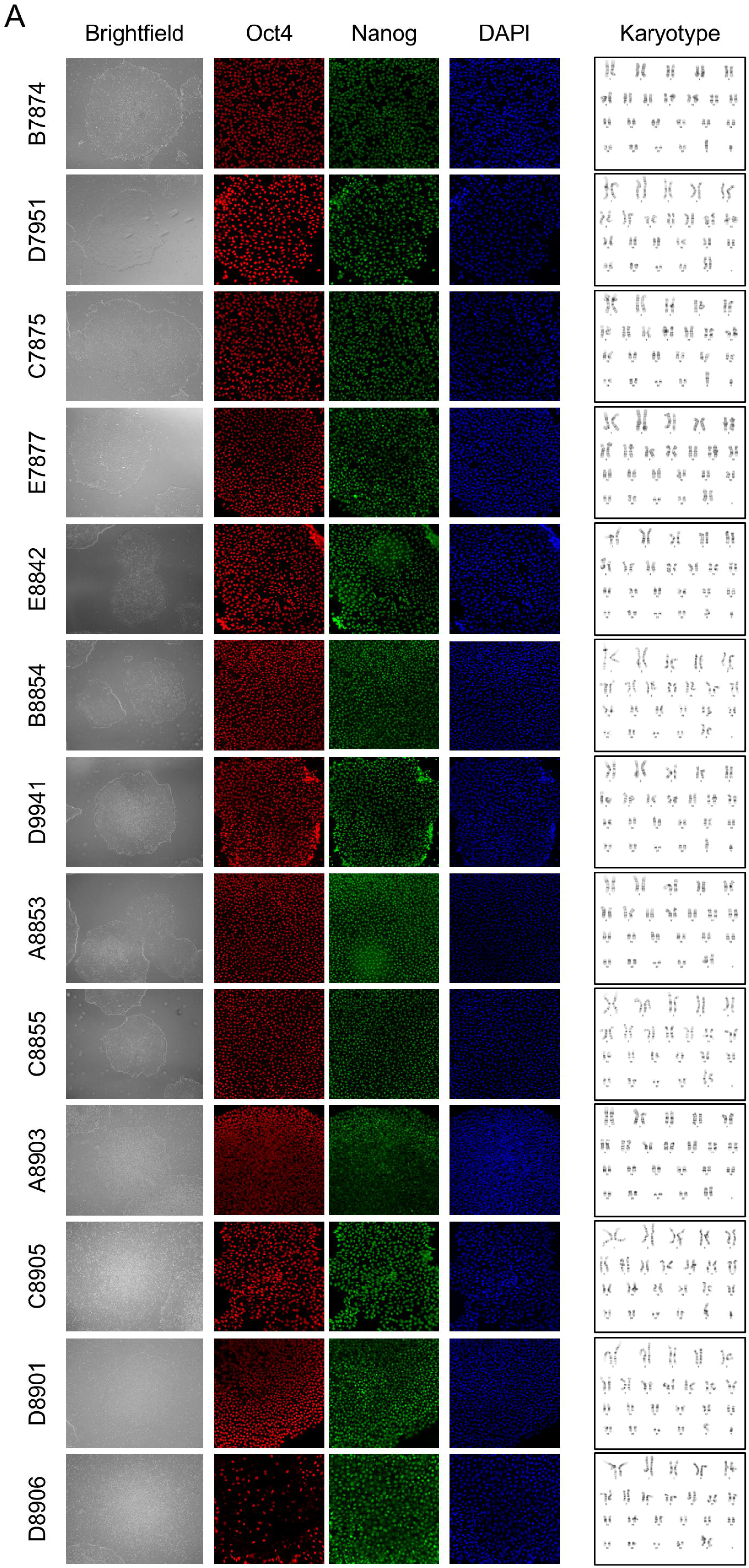
Patient cell lines pluripotency quality control. A) The pluripotent cell lines established for this work, derived from healthy controls and patients with MS, were tested for pluripotency and underwent quality control analysis. Additional cell lines were previously tested and characterized (Daviaud et al., 2023). Brightfield microscopy was used to assess IPSCs morphology and general health. Oct4 and Nanog immunofluorescence counterstained with DAPI was performed to verify IPSCs pluripotency. Karyotype analysis was performed by G-band testing to confirm genetic stability after reprogramming.

### Generation of neural precursor cells

Neural precursor cells (NPCs) were produced following a protocol previously described (Gunhanlar et al., 2018) with minor modification. Briefly, iPSCs were dissociated with EDTA (0.5 mM, Millipore) for 5-6 minutes at 37°C. Embryonic bodies (EBs) were generated by transferring 4000 cells per well of an ultra-low attachment 96 well plates in mTeSR Plus supplemented with Thiazovivin (2 µM, Millipore). After 2 days, the medium was changed to neural induction media (StemDiff, STEMCELL) and EBs were cultured for another 4 days. On day 7, EBs were slightly dissociated by mechanical trituration and cultured on Cultrex UltiMatrix (100 µg/ml, R&D systems) coated plates in neural induction medium (StemDiff, STEMCELL) for 7 days. At d15 the medium was switched to NPC medium consisting of DMEM/F12, 1% N2 supplement (ThermoFisher), 2% B27 without RA supplement (ThermoFisher), 20 ng/ml epithelial growth factor (Peprotech), 10 ng/ml basic fibroblast growth factor (Peprotech) and 1% penicillin/streptomycin (ThermoFisher). At d15, cells were considered pre-NPCs and could be passaged and cryopreserved at confluence. From passage 3, cells were considered NPCs and were used for histologic analysis and for neural differentiation.

Neural differentiation was achieved by mitogen withdrawal. NPCs were cultured for 10 days on Cultrex coated plates with differentiation media consisting of DMEM/F12, 1% N2 supplement and 2% B27 with RA supplement (ThermoFisher) and were then fixed and analyzed by immunostaining.

### Generation of human cerebral organoids

C-organoids were generated from human iPSCs and processed for analysis as described (Lancaster and Knoblich, 2014; Daviaud et al., 2023) with minor modifications. iPSCs were washed with Dulbecco’s phosphate-buffered saline (DPBS, ThermoFisher) and dissociated with EDTA 0.5 mM (Millipore). A total of 8 x 10^3^ cells were seeded into each well of an ultra-low attachment 96-well plate (Corning) to form embryoid bodies (EBs) in mTeSR Plus medium supplemented with 4µM of Thiazovivin (Millipore) for the first 2 days. The medium was changed every other day to the same medium without Thiazovivin for another 2-3 days. After 4-5 days of culture or when EBs reached ∼500-600 µm in diameter and the surface tissue began to brighten, EBs were cultured in neural induction medium (StemDiff, STEMCELL). After neuroepithelium emergence (typically at ∼ day 9-10), embryoid bodies were embedded in 15 µL Matrigel droplets and cultured in 6 well plates containing c-organoid differentiation medium consisting of 1:1 DMEM-F12 and Neurobasal medium (Gibco), with addition of 0.5% N2 supplement (Life Technologies), 0.5% ml MEM-NEAA (Gibco), 1% Glutamax (Gibco), 1% B27 supplement without vitamin A (Life Technologies), 0.1 µM of 2-mercaptoethanol (Millipore), 2.6 µg/ml insulin (Sigma Aldrich) in static culture for 4 days. Organoids were cultured in c-organoid differentiation medium supplemented with vitamin A on an orbital shaker (CO2 Resistant Shakers, ThermoFisher) at 80 rpm. Organoids were cultured up to 200 days while radial growth and neuroepithelial bud formation was observed.

### Generation of human spinal cord organoids

C-organoids were generated from human iPSCs and processed for analysis as described elsewhere (Lee et al., 2022; Xue et al., 2023) with minor modifications. iPSCs were washed with Dulbecco’s phosphate-buffered saline (DPBS, ThermoFisher) and dissociated with EDTA 0.5 mM (Millipore). A total of 8 x 10^3^ cells were seeded into each well of an ultra-low attachment 96-well plate (Corning) to form embryoid bodies (EBs) in mTeSR Plus medium supplemented with 4µM of Thiazovivin (Millipore) for the first 2 days. On day 3 to totality of the culture media was removed and replaced with a differentiation media consisting of 1:1 DMEM-F12 and Neurobasal medium (Gibco), with addition of 0.5% N2 supplement (Life Technologies), 0.5% ml MEM-NEAA (Gibco), 1% Glutamax (Gibco), 1% B27 supplement without vitamin A (Life Technologies), 0.1 µM of 2-mercaptoethanol (Millipore), 2.6 µg/ml insulin (Sigma Aldrich) supplemented with 10µM of SB431542 (STEMCELL), 2 µM of CHIR99021 (STEMCELL) and 0.5 µM of LDN-193189 (STEMCELL). Organoids were kept in culture for 8 days with half media change every other day. On day 11, organoids were embedded in 15 µL Matrigel droplets and cultured in 6 well plates containing differentiation medium supplemented with vitamin A and 15 ng/ml of BMP4 (PEPROTECH) on an orbital shaker (CO2 Resistant Shakers, ThermoFisher) at 80 rpm. On day 16, media was changed to a differentiation media supplemented with 10 ng/ml BDNF and 10 ng/ml GDNF (PEPROTECH). Organoids were cultured for 64 days.

### Histological Analysis

At D42 and D120 to D200 of culture, c-organoids were washed in D-PBS and fixed in 4% PFA (ThermoFisher) for 20 minutes at 4°C. After three washes with D-PBS, the c-organoids were cryoprotected in 30% sucrose overnight at 4°C followed by snap freezing in OCT compound (ThermoFisher) and stored at −20°C. Cryosections of organoids were cut at 15 µm thickness using a cryostat (Leica CM 1950) and mounted on microscope slides (Histobond^+^, VWR).

For immunofluorescence, slides were thawed to room temperature before being outlined with a PAP pen (Millipore) to create a hydrophobic barrier. Slides were washed and permeabilized with PBS supplemented with 0.1% Triton X-100 (Millipore Sigma). Non-specific binding sites were blocked with PBS supplemented with 0.1% Tween 20, 4% Bovine Serum Albumin (ThermoFisher) and 10% Normal Goat Serum (ThermoFisher) for 1 hour at RT. Slides were then incubated overnight at 4°C with the following primary antibodies diluted in blocking solution: mouse anti-APC (1:100, Calbiochem), mouse anti-ChAT (1:200, ThermoFisher), rat anti-CTIP2 (1:500, Abcam), guinea pig anti-DCX (1:500, Millipore), Rabbit anti-E2F1 (1:400, ThermoFisher), mouse anti-EOMES (1:100, ThermoFisher), mouse anti-GAD67 (1:200, Abcam), mouse anti-GFAP (1:500, Novus Biologicals), mouse anti-Ki67 (1:400, Millipore), chicken anti-MBP (1:500, ThermoFisher), mouse anti-Nanog (1:400, Abcam), mouse anti-O4 (1:200, R&D systems), rabbit anti-Oct4 (1:400, Abcam), rabbit anti-Olig2 (1:200, Abcam), rabbit anti-p16 (1:200, ThermoFisher), rabbit anti-p21 (1:200, ThermoFisher), rabbit anti-p27 (1:400, ThermoFisher), rabbit anti-p57 (1:400, ThermoFisher), mouse anti-Pax6 (1:100, Abcam), rabbit anti-PAK1 (1:100, ThermoFisher), mouse anti-SOX2 (1:400, Abcam), rabbit anti-TBR1 (1:400, Abcam), rabbit anti-vGluT1 (1:200, Abcam). After washing, slides were incubated with appropriate Alexa-coupled secondary antibodies (ThermoFisher) diluted in blocking solution for 1 hour at RT and counterstained with DAPI, before mounting with Fluoromount Aqueous Mounting Medium (Millipore).

Immunofluorescence images were collected using a fluorescence microscope (Zeiss Imager M2) or a confocal fluorescence microscope (Zeiss LSM 510) and processed using Zen software and Fiji software (Schindelin et al., 2012).

### Experimental design and statistical analysis

C-organoids were derived from 20 different patient cell lines, with a minimum of 4 cell lines per subtype of MS. Three different subclone were generated for each cell line. For each cell line, 3 to 4 independent batches were generated, with 2 to 3 organoids per batch. Subsequently, 1-3 representative images from each organoid were quantified (Fiji software) (Schindelin et al., 2012).

To determine the percentage of stained cells in the c-organoid sections, the appropriate immunohistochemical markers counterstained with DAPI were used for quantification. For each section, regions of interest were generated in 250 µm (width) x 300 µm (height) radial columns spanning all cortical layers near the organoid surface to normalize the analyzed area.

Statistical analysis was performed using GraphPad Prism 8. Differences between multiple conditions were determined using a one-way analysis of variance (ANOVA) test. If the ANOVA residuals were normally distributed (Shapiro-Wilk test), a Tukey’s post hoc multiple comparison test was performed. In the case that the ANOVA residuals were not normally distributed (Shapiro-Wilk test), a Kruskal-Wallis test followed by Dunn’s multiple comparison test was performed. A chi-squared test was used to analyze frequency distributions.

All data are presented as the mean value with standard error of the mean (SEM) unless otherwise noted. Results were considered significant if p < 0.05.

## RESULTS

### Cerebral organoid derived from patients with MS develop and mature over time

Multiple sclerosis is a heterogeneous disease whose course and severity can vary greatly from patient to patient. To best capture the diversity/variability of the disease, we decided to generate IPS cell lines derived from 10 male and 10 female subjects (Table 1), including 6 healthy controls and 14 MS patients. The age of the patients ranged from 23 to 67 years old with no significant differences between healthy controls and the different MS groups (One-Way ANOVA, p = 0.3450) (Table 2).

Each patient IPS cell line exhibited a normal phenotype in culture. Expression of pluripotency markers, such as Oct4 and Nanog were confirmed by immunofluorescence and each cell line had a normal Karyotype (Figure 1).

Cerebral organoids were generated using a previously described protocol (Lancaster and Knoblich, 2014; Daviaud et al., 2023) (Figure 2A). After 40-50 days in vitro, c-organoids exhibited immature cortical structures consisting of the ventricle aligned with proliferating cells, surrounded by the ventricular zone (VZ) containing the SOX2^+^ stem cell pool, the subventricular zone (SVZ) containing TBR2^+^ intermediate progenitors and DCX^+^ neuroblasts, and the cortical plate (CP) containing mature neurons (Figure 2B and 2D). After 120-150 days in vitro, c-organoids continued to mature, and exhibited more mature neurons, such as GABAergic and Glutamatergic neurons, but also mature myelinating oligodendrocytes and astrocytes (Figure 2C and 2E).

**Figure 2:**
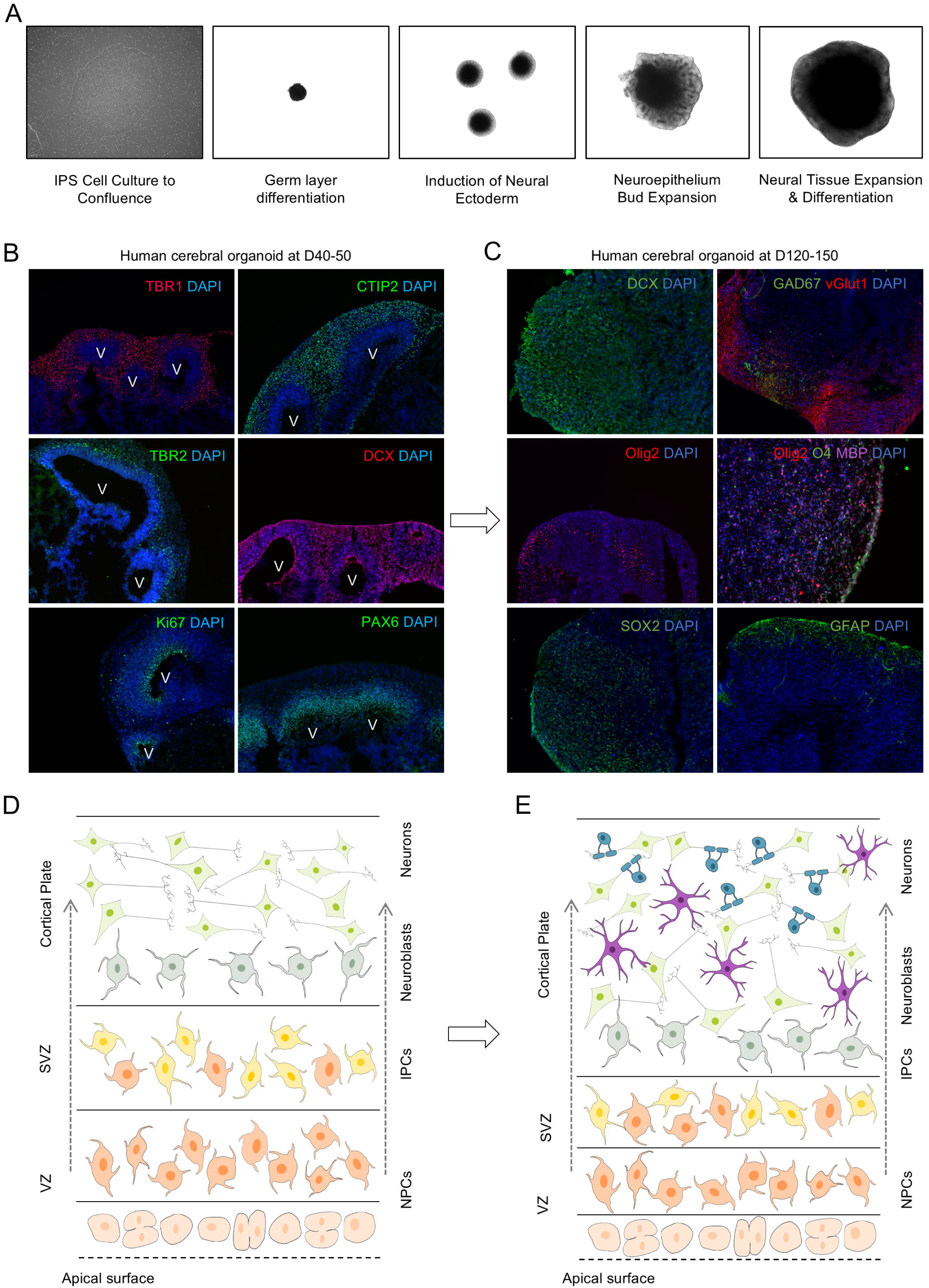
Cerebral organoids derived from patients with MS develop and mature over time. A) Pictures of IPS cells derivation into cerebral organoid using brightfield microscopy. B) Immunofluorescence of human cerebral organoids at D40-50 for the major cell types detected: Ki67^+^ proliferating cells, SOX2^+^ neural precursors, TBR2^+^ intermediate progenitors, DCX^+^ neuroblasts, TBR1^+^ and TBR2^+^ mature cortical neurons. V: ventricles. C) Immunofluorescence of human cerebral organoids at D120-150 for SOX2^+^ neural precursors, GFAP^+^ astrocytes, Olig2^+^ oligodendrocytic cells, DCX^+^ neuroblasts, GAD67^+^ and vGlut1^+^ GABAergic and glutamatergic neurons respectively and Olig2^+^O4^+^MBP^+^ myelinating oligodendrocytes. D) Schematic representation of the cortical structures found in cerebral organoids at D40-50 including the ventricular zone containing the stem cell pool and most proliferating cells, the subventricular zone (SVZ) containing mostly IPCs, the outer SVZ (oSVZ) and the cortical plate (CP) containing neuroblasts and mature neurons. E) Schematic representation of the structures found in cerebral organoids at D120-150. At this point there is no clear distinction between cortical layers. The rim of the organoids contains stem cells but mostly mature neurons, astrocytes, and myelinating oligodendrocytes.

### CDKi involved in MS organoids appears to be restricted to p21

CDKIs are proteins that bind to and inhibit the activity of CDKs, thereby controlling the cell cycle. Two major classes of CDK inhibitors have been identified. The INKs family includes p15, p16, p18 and p19 which bind to and inhibit the activities of CDK4 and CDK6. The CIP/Kips family includes p21, p27, p28 and p57 which can bind to and inhibit the activities of a wide range of CDK-cyclin complexes (Besson et al., 2008) (Figure 3A).

**Figure 3:**
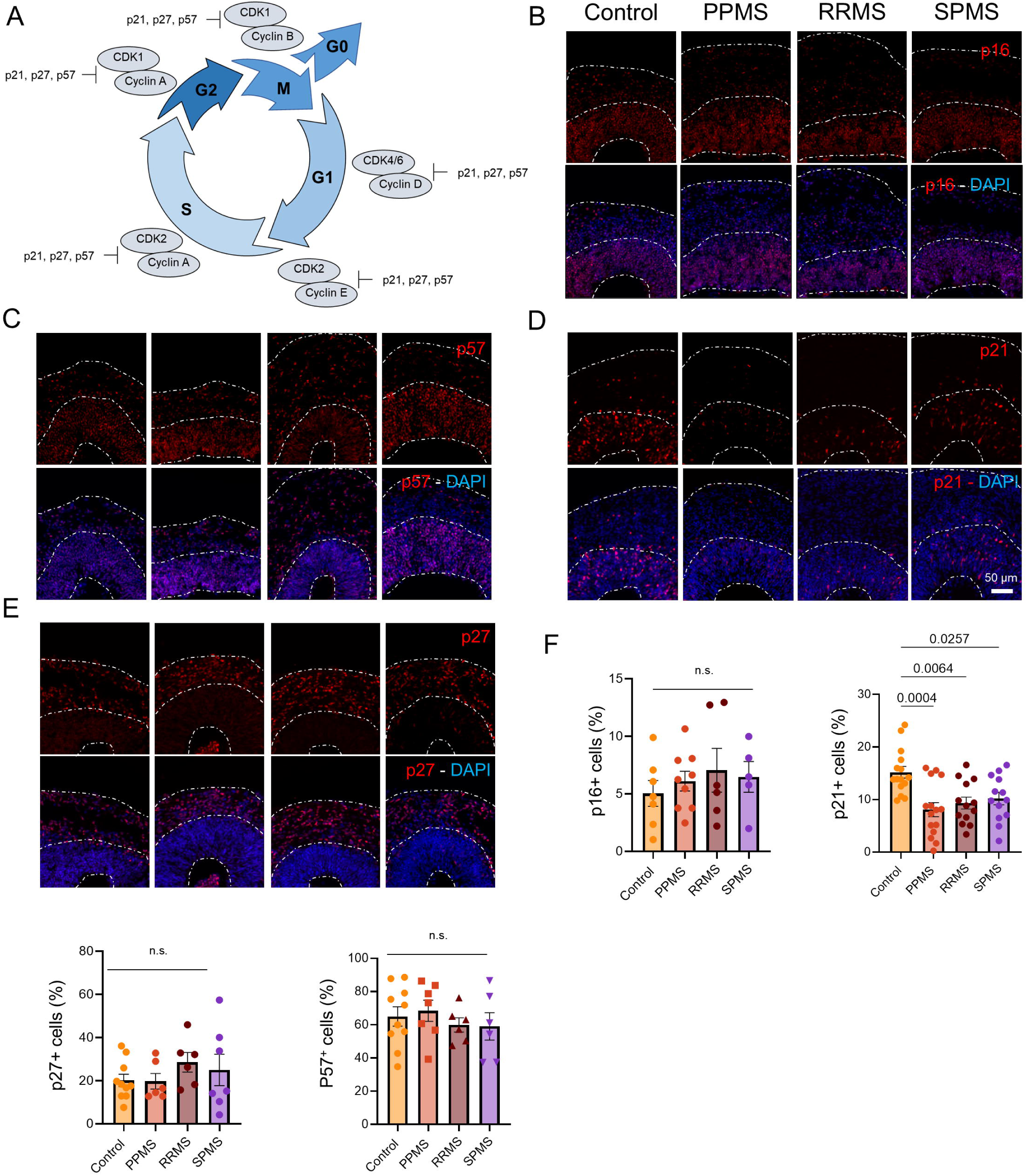
CDKi involved in MS organoids appears to be restricted to p21. A) Schematic representation of the cell cycle and the involvement of the various CDK/cyclin complexes and the different CDK inhibitors. B) Representative images of an immunofluorescence against the CDKi p16 in c-organoids at D42 in the different conditions. No ectopic location or important change in p16 expression were observed. C) Representative images of an immunofluorescence against the CDKi p57 in c-organoids at D42 in the different conditions. p57 was evenly distributed among the layers of c-organoids. No ectopic location or important change in p57 expression were observed. D) Representative images of an immunofluorescence against the CDKi p21 in c-organoids at D42 in the different conditions. p21 was mostly expressed in the lower layers (SVZ and VZ) of c-organoids. No ectopic location was detected, however, an important decrease of p21 expression was observed, especially in PPMS. E) Representative images of an immunofluorescence against the CDKi p27 in c-organoids at D42 in the different conditions. p27 was mostly expressed in the outer layers of c-organoids (CP). No ectopic location or important change in p27 expression were observed. F) Quantification for p16 (One-Way ANOVA, p = 0.7347), p27 (One-Way ANOVA, p = 0.5313), p57 (One-Way ANOVA, p = 0.7257) and p21 (One-Way ANOVA, p = 0.0004). To ensure quantification consistency, analyzed cortical area were cropped from the original picture to a 250 µm wide×300 µm image, spanning all cortical layers (VZ, SVZ, and CP). Each dot represents one organoid, 4-6 pictures were taken per organoid.

We previously highlighted the involvement of p21 in a c-organoid model of MS (Daviaud et al., 2023). However, we did not evaluate other CDKi expressions in MS organoids compared to controls. Indeed, p16 expression is increased in PPMS neural stem cells leading to senescence which could contribute to limited remyelination (Nicaise et al., 2019). p27 ensures cell cycle arrest and subsequent differentiation of oligodendrocyte precursor cells (OPCs) in mature oligodendrocytes (Casaccia-Bonnefil et al., 1997) and has also been described as a positive regulator of Schwann cell differentiation in vitro (Li et al., 2011). p57 regulates the number of divisions in OPCs before the onset of differentiation and can inhibit oligodendrocyte and Schwann cell differentiation. Therefore, all these CDKi may be important in MS pathogenesis.

We performed an immunofluorescence at d42 for p16, p21, p27 and p57 in organoids from patients with MS and healthy controls (Figure 3B-E). No ectopic location or significant expression difference was observed for p16, p27 and p57, while a decreased expression of p21 was found in MS organoids, especially in PPMS (Figure 3B-E). Quantification revealed no significant difference for p16 (One-Way ANOVA, p = 0.7347), p27 (One-Way ANOVA, p = 0.5313), p57 (One-Way ANOVA, p = 0.7257), while a significant decrease of p21 expression was found in MS organoids (One-Way ANOVA, p = 0.0004), particularly between control and PPMS (p = 0.0004) (Figure 3F).

### The CDKi involved in MS NPC in vitro appears to be limited to the cell cycle inhibitor p16

To confirm the results observed in cerebral organoids in a simpler model, we decided to perform immunofluorescence for the CDKi p16, p21, p27 and p57 on IPS-derived NPCs. We performed these analyses on 2 conditions: NPC in expansion and NPC after 10 days of neural differentiation (Figure 4).

**Figure 4:**
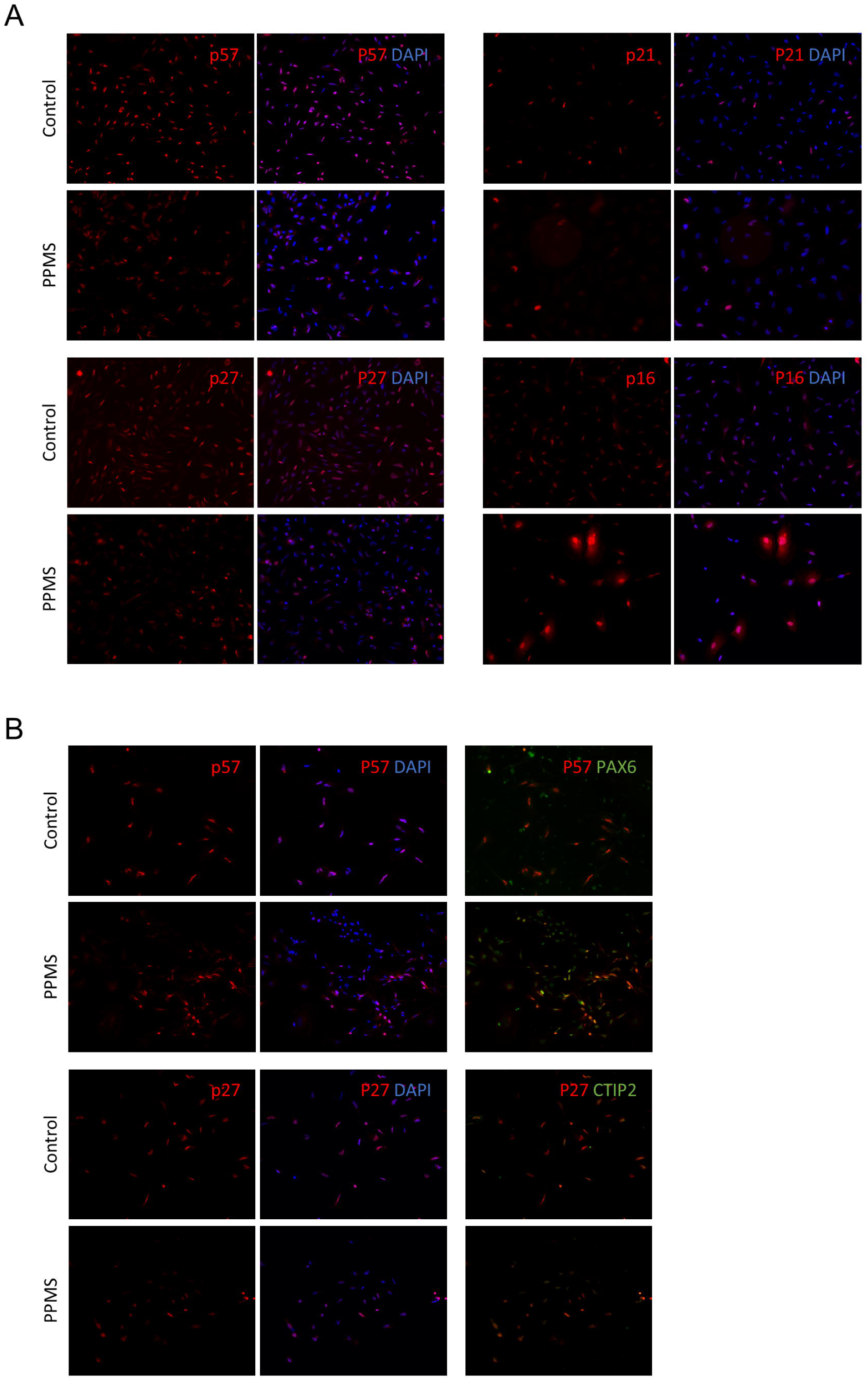
IPSC derived NPC and neural cell from PPMS exhibit p16 induced senescence. A) Representative images of an immunofluorescence performed on IPS-derived NPC in expansion for p57, p21, p27 and p16, in control and PPMS samples. Only p16 expression was different between the two conditions. B) Representative images of an immunofluorescence performed on IPS-derived NPC, 8 days after mitogens withdrawal for p57 and p27 in control and PPMS samples. p57 and p27 were also co-stained with PAX6 and CTIP2, respectively.

In proliferating PPMS NPCs, no significant change in p21 or p27 expression was observed compared to control. However, a significant increase in p16 expression was detected, confirming previous results (Nicaise et al., 2019). Furthermore, a possible translocation of p57 from the nucleus to the cytoplasm was detected in PPMS NPCs compared to control (Figure 4A).

In contrast no significant change in p27 and p57 expressions were detected in differentiated NPCs in PPMS compared to control (Figure 4B).

### P21 regulators E2F1 and PAK1 are involved in MS pathogenesis

The CDKi p21 is significantly decreased in MS organoids, especially in PPMS. In addition, the functions of p21 in oligodendrocyte differentiation make it an interesting protein to study for MS. For these reasons, we analyzed two proteins, PAK1 and E2F1, as they are both involved in p21 regulation (Hiyama et al., 1998; Gartel et al., 2000; Wang et al., 2018), and also in oligodendrocyte maturation and myelination capacity (Figure 5A) (Thurnherr et al., 2006; Magri et al., 2014; Wu et al., 2022).

**Figure 5:**
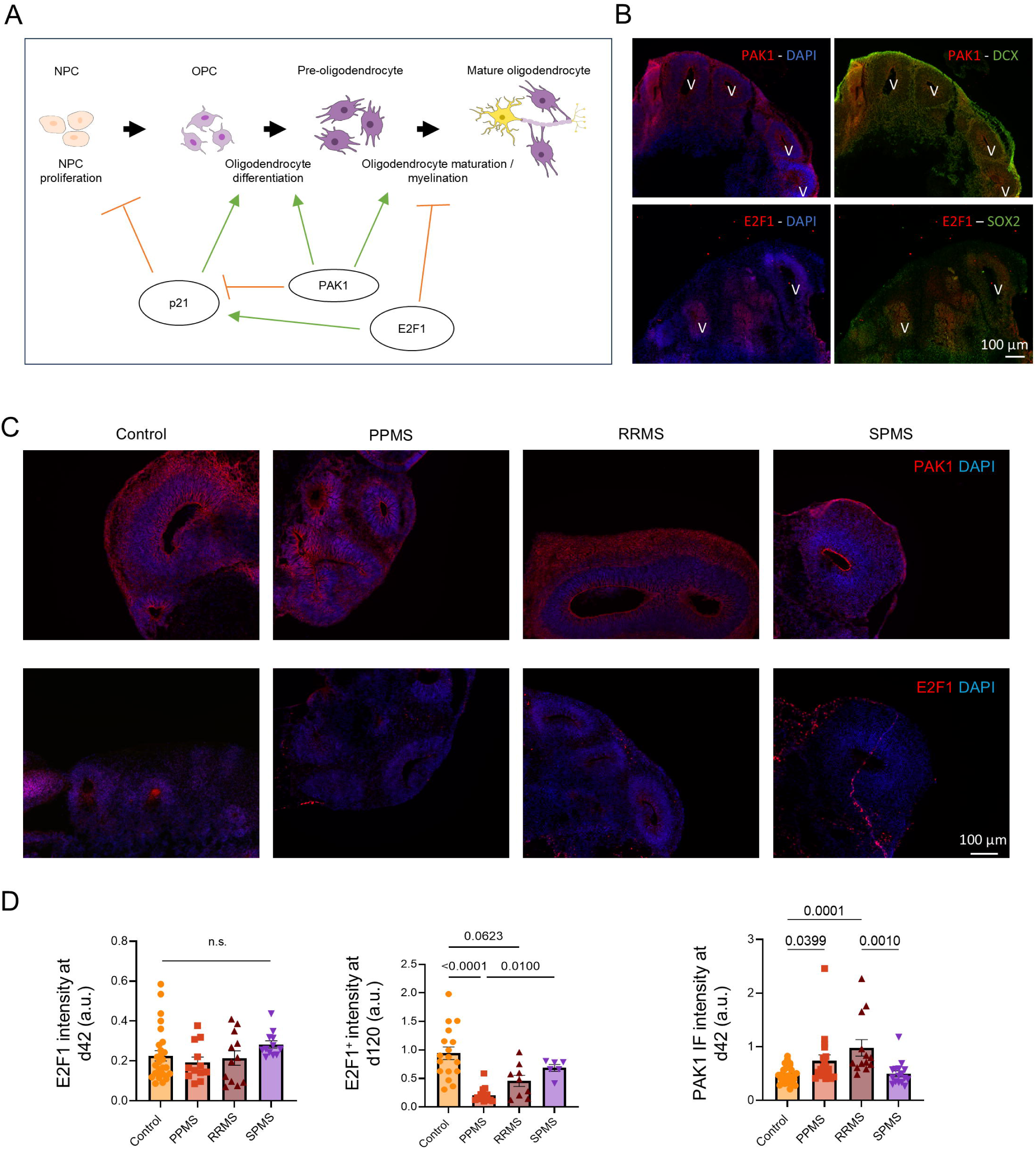
P21 regulators E2F1 and PAK1 are involved in MS pathogenesis. A) Schematic representation of the involvement of PAK1 and E2F1 in p21 regulation, but also in oligodendrocyte differentiation, maturation, and myelination capacity. B) Immunofluorescence for PAK1 co-stained with neuroblast marker DCX (Top) and E2F1 co-stained with stem cell marker SOX2. C) Representative images of an immunofluorescence for PAK1 and E2F1 in organoids at D42 in control, PPMS, RRMS and SPMS samples. D) Quantifications for E2F1 in c-organoids at D42 (Kruskal-Wallis, p = 0.0898), for E2F1 in c-organoids at D120 (Kruskal-Wallis, p < 0.0001) and for PAK1 at D42 (Kruskal-Wallis, p < 0.0001). Each dot represents an organoid generated from different batches and patient cell lines. 4-6 images were taken per organoid.

We first determined the localization of E2F1 and PAK1 expression in c-organoids at d42 by immunofluorescence. PAK1 was mainly expressed in the outer layers of the organoid cortical structure, such as the cortical plate, suggesting that PAK1 was mainly expressed in mature cells, such as neuroblasts, astrocytes or oligodendrocytes (Figure 5B). E2F1 was mainly expressed in the lower layer of the cortical structure, especially in the ventricular zone, which indicates that E2F1 was mainly expressed in the progenitor cells, such as NPCs (Figure 5B).

We then performed an immunofluorescence for PAK1 and E2F1 in cerebral organoids from patients with MS compared to organoids from healthy controls (Figure 5C). Quantification showed no significant difference in E2F1 expression between the different conditions at D42 (Kruskal-Wallis, p = 0.0898). At D120 a significant decrease of E2F1 expression was observed in PPMS organoids compared to control but also to SPMS (Kruskal-Wallis, p < 0.0001) while no difference was noted with RRMS. At D42, an increase of PAK1 expression was also detected in MS organoids compared to control, especially PPMS and RRMS (Kruskal-Wallis, p < 0.0001) (Figure 5D).

These results suggest that the dysregulation of p21 may be induced or associated with changes in E2F1 and PAK1 expression in the cerebral organoid model of MS.

### Oligodendrocyte maturation and myelination capacity is reduced in MS organoids

The differentiation of NPCs into mature oligodendrocytes is a complex process. First, NPCs must differentiate into NG2^+^ OPCs, then mature into O4^+^ pre-oligodendrocytes, which mature into GALC^+^/MBP^+^ myelinating oligodendrocytes. During each of these steps, oligodendroglial cells express Olig2 (Figure 6A).

**Figure 6:**
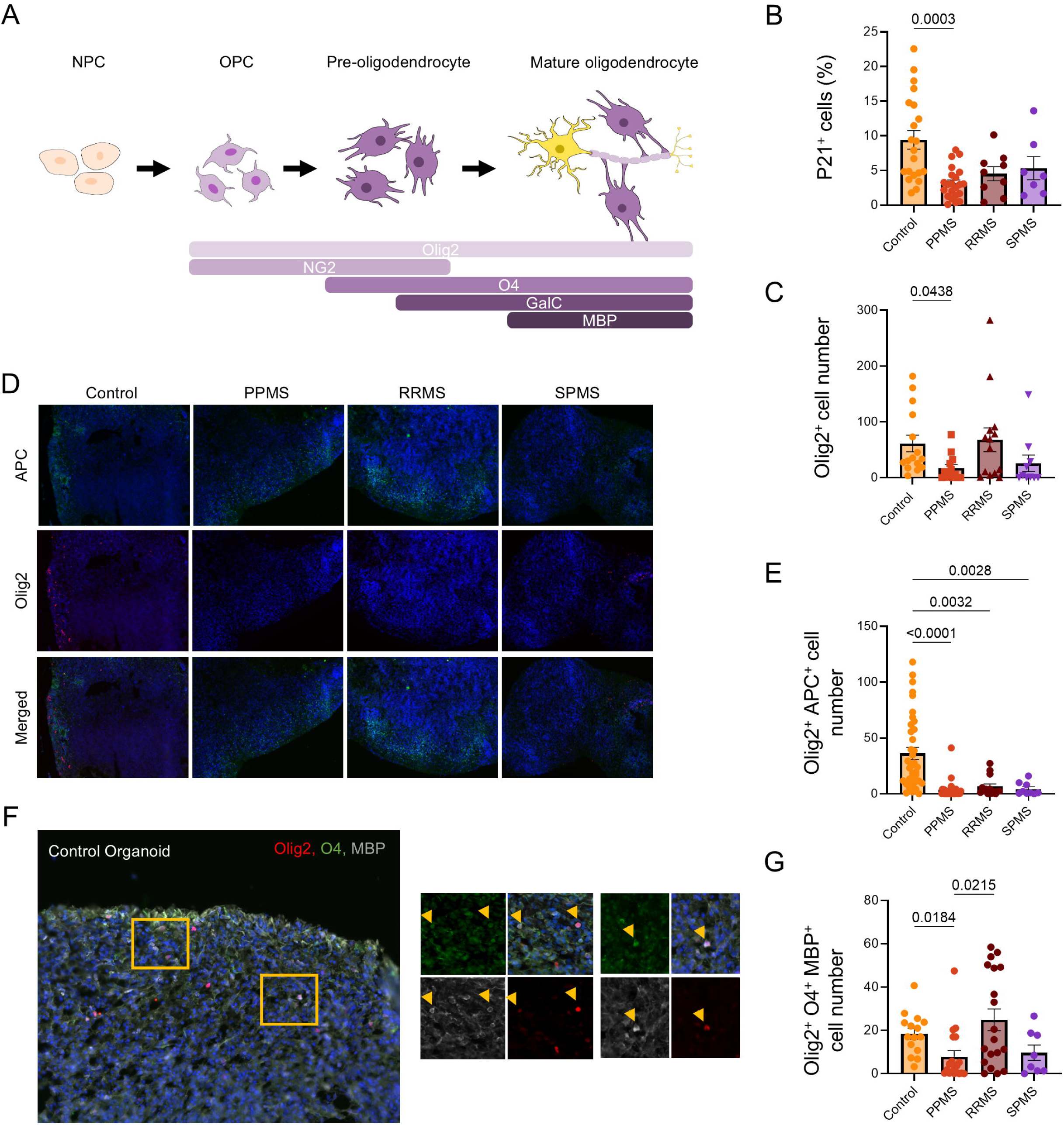
Oligodendrocyte maturation and myelination capacity is reduced in MS organoids at D120. A) Schematic representation of the different markers expressed during the differentiation of NPCs into OPCs and their maturation into pre-oligodendrocytes and mature myelinating oligodendrocytes. B) Quantification of p21^+^ cells in c-organoids at D120 highlighting a significant change in PPMS organoids (Kruskal-Wallis, p = 0.0009). C) Quantification of Olig2^+^ cells in c-organoids at D120, showing a significant difference in PPMS organoids (Kruskal-Wallis, p = 0.0109) D) Representative images of an immunofluorescence for Olig2 and APC, a marker of astrocytes and mature oligodendrocytes. Olig2^+^APC^+^ double positive cells indicate mature oligodendrocytes. E) Quantification of Olig2^+^APC^+^ double positive cells in c-organoids at D120. A significant difference was detected (Kruskal-Wallis, p = <0.0001). F) Representative images of an immunofluorescence for Olig2, O4 and MBP in control c-organoids at D120. The orange squares represent the zone of interest highlighted on the right panel. Orange arrowheads indicate triple positive cells. Colocalization of those three markers indicates the presence of myelinating oligodendrocytes. G) Quantification of Olig2^+^O4^+^MBP^+^ triple positive cells in c-organoids at D120. A significant difference was detected (Kruskal-Wallis, p = 0.0057). Each dot represents an organoid generated from different batches and patient cell lines. 4-6 images were taken per organoid.

In our previous work, we highlighted a decrease of oligodendroglial cell marker Olig2 in MS organoids compared to control at D42. At this time point, few O4^+^ cells were detected in control, while no positive cells were found in PPMS organoids, and no myelin could be detected. To further study oligodendrocyte differentiation and maturation in MS organoids, we decided to keep our organoids in culture for 120 to 200 days in vitro. By this time point, organoids are more mature and exhibit not only GABAergic and glutamatergic neurons, but also mature myelinating oligodendrocytes and astrocytes (Figure 2C and 2E).

To assess its expression at D120, we performed an immunofluorescence for p21 in our organoid model of MS. Quantification showed a significant decrease in p21 expression in PPMS organoids compared to control (Kruskal-Wallis, p = 0.0009) (Figure 6B). This result indicates that the p21 decrease in MS organoids occurs continuously from D28 to D120 and is not a transient effect.

To examine oligodendrocyte maturation and myelination capacity in our long-term cultured organoids, we studied Olig2 expression in oligodendroglial cell population. Quantification using immunofluorescence revealed a significant decrease of Olig2^+^ cell number in PPMS compared to control. Interestingly a slight decrease in Olig2^+^ cells was also observed in SPMS samples but didn’t reach significance (Kruskal-Wallis, p = 0.0109) (Figure 6C).

To further assess oligodendrocyte maturation, we performed a double immunofluorescence for the oligodendroglial cell marker Olig2 and the mature oligodendrocyte marker APC (Figure 6D). Double positive cells indicated the presence of mature oligodendrocytes in our organoid model of MS. Quantification indicated a significantly lower number of Olig2^+^/APC^+^ cells in PPMS, RRMS and SPMS organoids compared to control (Kruskal-Wallis, p < 0.0001) (Figure 6E).

We also analyzed the maturation of oligodendrocytes into myelinating oligodendrocytes in our c-organoids, by triple immunofluorescence for Olig2, the mature oligodendrocyte marker O4, and the myelinating oligodendrocyte marker MBP (Figure 6F). Triple positive cells were found in control organoids, while lower numbers were found in MS organoids. Quantification highlighted a significantly lower number of Olig2^+^ O4^+^ MBP^+^ cells in PPMS, with a slight reduction in SPMS organoids compared to control (Kruskal-Wallis, p = 0.0057), while no difference was found between control and RRMS (Figure 6G).

Finally, we performed an immunofluorescence for the myelin marker MBP and the neuronal marker DCX to verify the quality of the myelin produced in our organoid at d120-150. In control organoids we could observe colocalization of the neuronal marker DCX and the myelin marker MBP, highlighting formation of myelin around the axons, whereas little to no colocalization was observed in PPMS organoids (Figure 7A). We used confocal microscopy to further confirm the quality of myelin in control organoids (Figure 7B) and performed 3D reconstruction of stacked images (Figure 7C). In control organoids, we observed perfect colocalization of the neuronal marker DCX and the myelin marker BMP. The colocalization was found along the axons, highlighting a proper myelination of axons in our organoid model at this time-point.

**Figure 7:**
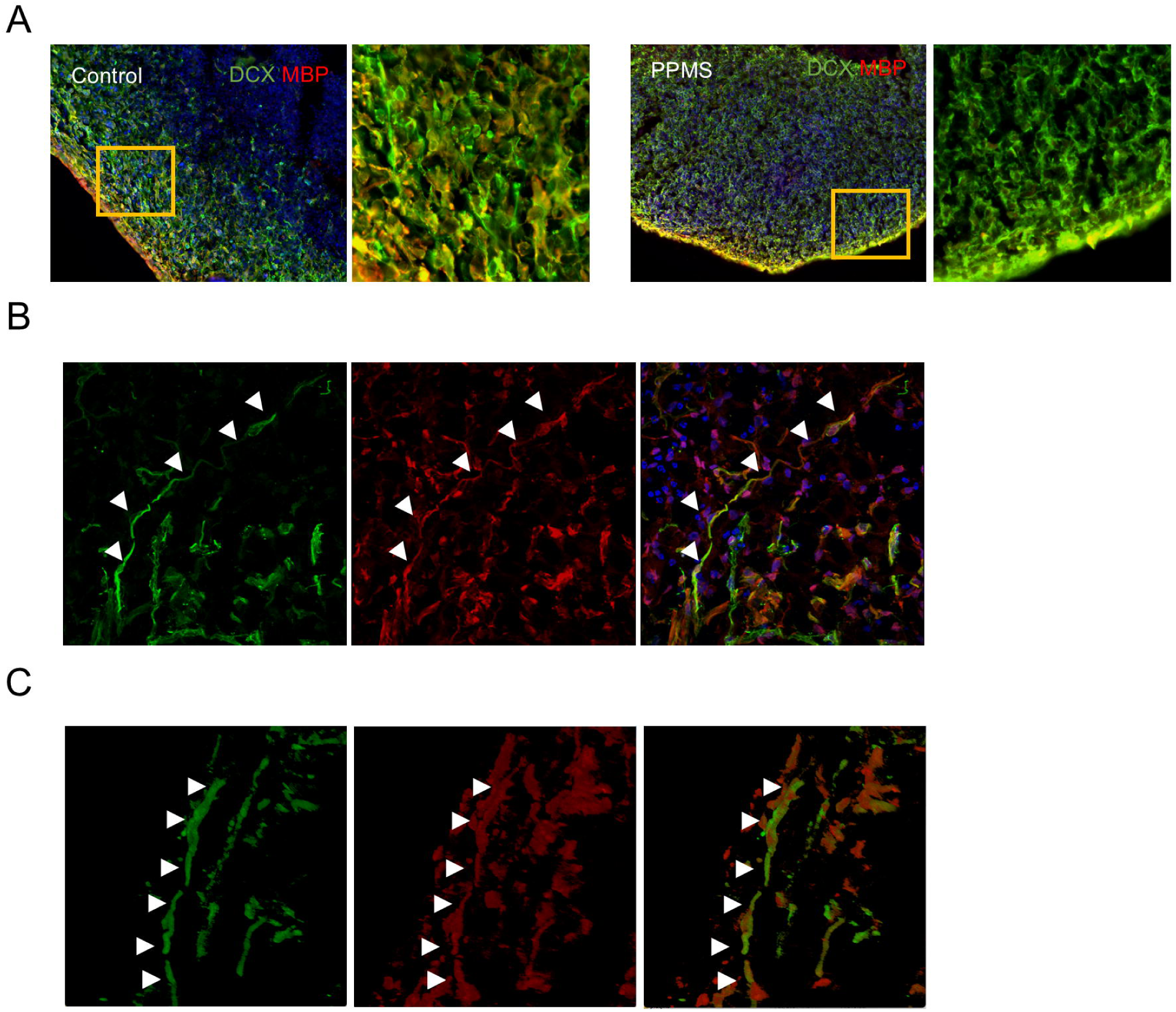
Organoid model of MS contains myelinating neurons. A) Representative images of an immunofluorescence for neuron marker DCX and myelin marker MBP in control and PPMS organoids at D120. The orange squares indicate a zone of interest magnified on the right panel. B) Confocal microscopy was performed to confirm colocalization of axons with myelin marker, showing proper myelination. White arrowhead indicates DCX and MBP colocalization. C) 3D reconstruction of stacked images taken by confocal microscopy to further observed myelinated axons in c-organoids. White arrowhead indicates DCX and MBP colocalization.

Taken together, these results suggest that a defect in oligodendrocyte differentiation and maturation occurs early during organoid maturation and remains unchanged after long-term culture, resulting in a defect of myelination, which may partially explain the clinical phenotype of patients, especially of those with PPMS. Moreover, these results correlate in time with a corresponding decrease in p21 expression.

### Neurons and astrocyte differentiation are also affected in MS organoids

As stated before, at D120, organoids were more mature and exhibited GABAergic and glutamatergic neurons, mature myelinating oligodendrocytes and astrocytes (Figure 2C and 2E).

We studied astrocyte population as astrocytes may contribute to MS pathology by targeting dysregulated immune responses to the CNS and may lead to MS symptoms (Ponath et al., 2018). We identified astrocyte population by immunofluorescence for astrocyte marker GFAP (Figure 8A) and quantification showed a significant decrease of GFAP expression in MS organoids (Kruskal-Wallis, p = 0.0118) especially in PPMS samples compared to control and SPMS (Figure 8B).

**Figure 8:**
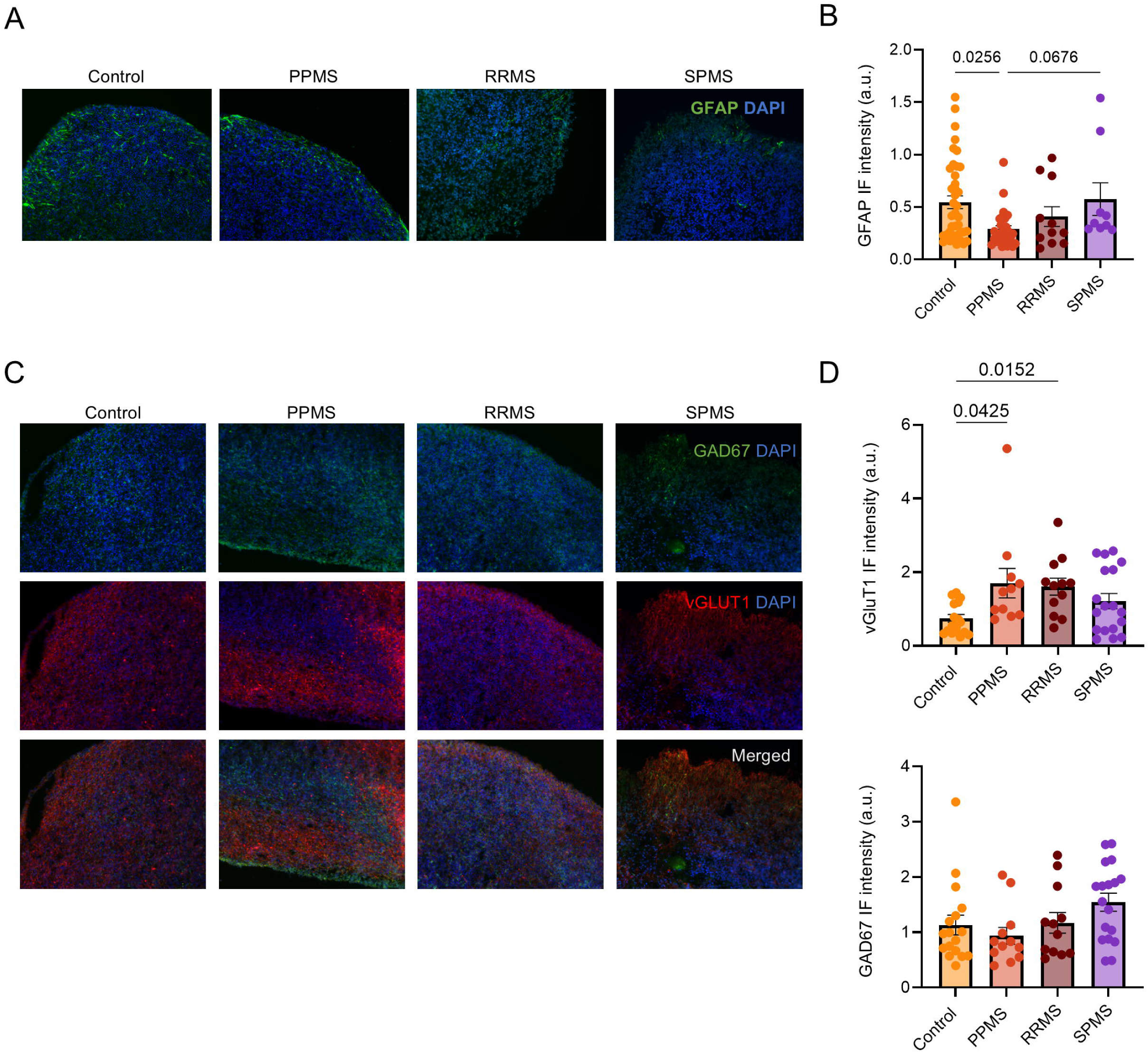
Neuronal and astrocytic differentiation are also affected in MS organoids. A) Representative images of an immunofluorescence for the astrocyte marker GFAP in c-organoids at D120. B) Quantification of GFAP fluorescence intensity in c-organoids at D120. A significant difference was detected (Kruskal-Wallis, p = 0.0118). C) Representative images of an immunofluorescence for the GABAergic neuron marker GAD67 and the glutamatergic neuron marker vGlut1 in c-organoids at D120. D) Quantification of vGlut1 and GAD67 fluorescence intensity in c-organoids at D120. A significant difference was found for vGlut1 (Kruskal-Wallis, p = 0.0079) and GAD67 (Kruskal-Wallis, p = 0.0445).

We also examined GABAergic and glutamatergic neurons as a dysregulation of GABAergic/glutamatergic neurotransmission is associated with MS symptoms such as fatigue (Arm et al., 2021) or cognitive performance (Huiskamp et al., 2023). We performed an immunofluorescence against glutamatergic neuron marker vGlut1 and GABAergic neuron marker GAD67 (Figure 8C). Quantification revealed a significant increase of vGlut1 expression in PPMS and RRMS organoids compared to control (Kruskal-Wallis, p = 0.0079) (Figure 8D). No significant difference in GAD67 expression was found in MS organoids compared to control (Kruskal-Wallis, p = 0.0792) (Figure 8D).

### Spinal cord organoids derived from MS patient exhibit similar phenotype than c-organoids

Spinal cord abnormalities are common in MS and include a variety of pathological processes, such as inflammatory demyelination, neuroaxonal loss and gliosis. Ultimately these changes result in motor weakness associated with gait difficulties, sensory disturbances, as well as bladder and bowel dysfunction (Walker, 2000).

We created spinal cord organoids (SCO) from a minimum of 3 healthy controls as well as 9 patients with MS to have a more comprehensive review of MS pathogenesis and development.

We first assessed mature neuron populations in our spinal cord organoid model of MS (Figure 9A). We performed an immunofluorescence for neuroblast marker DCX, but also glutamatergic neuron marker vGlut1 and GABAergic marker GAD67 (Figure 9A). Quantification revealed an increase of neurogenesis in SCO from MS patients, especially in its progressive forms, but also a significant higher expression of GABAergic neurons (Kruskal-Wallis, p = 0.0170), while no significant difference was detected for glutamatergic neurons (Kruskal-Wallis, p = 0.4578) in MS SCO compared to control (Figure 9B). Similarly to our work on cerebral organoids, we detected in our patient derived SCO a significant dysregulation of the excitatory/inhibitory neurons balance. An analysis of ChAT^+^ motor neurons did not detect any difference in ChAT^+^ motoneurons expression in MS SCO compared to control (data not shown).

**Figure 9:**
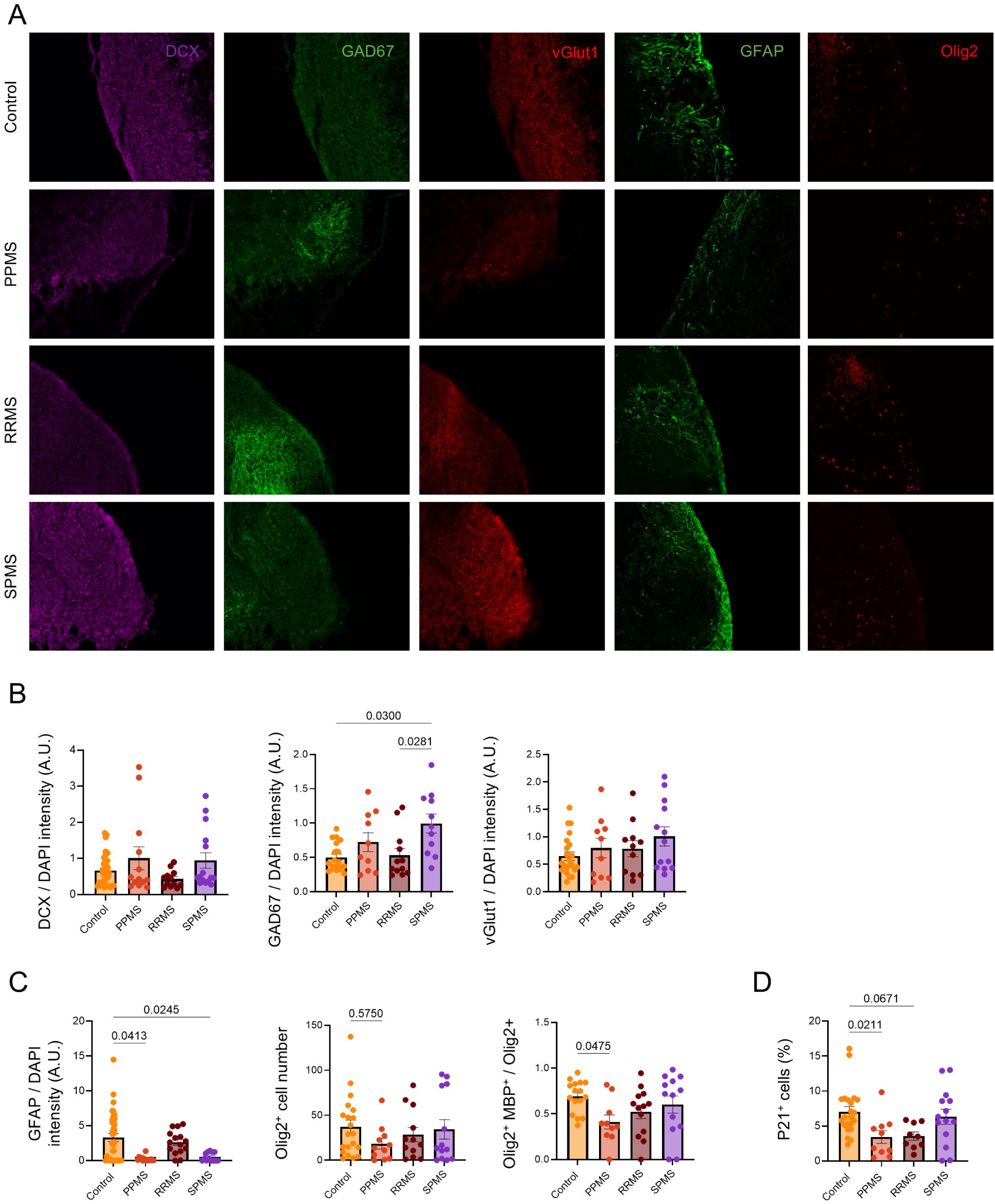
Spinal cord organoids derived from MS patient exhibit similar phenotype than c-organoids. A) Representative images of immunofluorescence for neuroblast marker DCX, GABAergic neuron marker GAD67, glutamatergic neuron marker vGlut1, astrocyte marker GFAP and oligodendrocyte marker Olig2 in spinal cord organoids at D64. B) Quantification for DCX, vGlut1 and GAD67 fluorescence intensity in spinal cord organoids at D64. Quantification showed no significant difference in DCX expression (Kruskal-Wallis, p = 0.2973) and vGlut1 (Kruskal-Wallis, p = 0.4578), while a significant difference was detected in GAD67 (Kruskal-Wallis, p = 0.0170) expression in MS SCO compared to control. C) Quantification for GFAP fluorescence intensity, Olig2^+^ cell number and for Olig2^+^O4^+^MBP^+^ myelinating oligodendrocyte number in spinal cord organoids at D64. Quantifications revealed a significant difference in astrocyte population (Kruskal-Wallis, p = 0.0013). No significant difference was observed for Olig2^+^ cells (Kruskal-Wallis, p = 0.4096). A significant reduction in myelinating oligodendrocyte population (One-way ANOVA, p = 0.05) expression in PPMS SCO compared to control. D) Quantification for p21^+^ cells showed a significant decrease in p21 expression (Kruskal-Wallis, p = 0.0058) in PPMS SCOs compared to control while only a slight decrease for detected in RRMS samples.

We also analyzed glial cell populations by immunofluorescence for astrocyte marker GFAP and oligodendrocyte marker Olig2 (Figure 9A). Quantifications revealed a significant decrease in astrocytic cells in MS organoids compared to control (Kruskal-Wallis, p = 0.0013) particularly in PPMS and SPMS. Only a slight decrease in Olig2^+^ cells was observed in PPMS organoids (Figure 9C). To assess oligodendrocyte myelination capacity in our SCO model of MS we performed an immunofluorescence for Olig2, O4 and MBP. A significant decrease in myelinating oligodendrocyte population was detected in our SCO model of MS (One-way ANOVA, p = 0.05) particularly in PPMS compared to control (Figure 9C).

Finally, we analyzed p21 expression (Figure 9D) and detected a decrease in PPMS and RRMS SCO (Kruskal-Wallis, p = 0.0058), confirming the involvement of p21 as observed in our cerebral organoids.

## DISCUSSION

We previously showed that c-organoids derived from patients with MS could be used as an innovative tool to better understand the genetic basis for phenotypic differences seen in MS. Using this model, we detected a decrease of proliferative capacity, notably PPMS, associated with a reduction of the progenitor pool and an increase of neurogenesis possibly due to an asymmetric shift of the mode cell division (Daviaud et al., 2023).

In this article we show that long-term cultured cerebral organoid from patients with MS exhibits a dysfunction of oligodendrocyte differentiation and maturation capacity leading to a decrease in myelination. In parallel, an increase of glutamatergic neurons associated with a decrease in astrocytic population was detected. These effects are linked to the dysregulation of the CDKi p21 and its regulators E2F1 and PAK1, while other CDKi, p16, p27 or p57, are not implicated in this process.

In this study, we developed and used 20 distinct patient-derived iPS cell lines and 3 clones were generated from each iPS cell line to enhance the reliability of our observations. IPSC cell lines were derived from 6 controls and 14 MS patients of different ages (from 23 to 67 years old), genders, and ethnicities, to ensure data validity. Statistical analysis confirmed there were no differences in age or gender distribution between the groups (Table 1-2). All iPS cell lines passed quality control and karyotype analysis, contributing to a robust and reproducible study design.

In our previous study we focused on precursor cells proliferation and differentiation capacity. Therefore, we used organoids at d42. At this time-point organoids contain a large pool of proliferating cells and neural stem cells, as well as intermediate progenitor cells, neuroblast and neurons but no mature astrocyte or oligodendrocyte could be observed (Figure 2B and 2D). However, long-term culture organoids further mature (Matsui et al., 2018), and generate oligodendrocytes and myelin after 20 to 30 weeks ((Renner et al., 2017; Madhavan et al., 2018) even responding to promyelinating drugs (Madhavan et al., 2018). Therefore, we decided to culture cerebral organoids up to day 120-150. At this time-point we observed mature neuronal populations such as GABAergic and glutamatergic neurons, mature glial cells such as astrocytes and myelinating oligodendrocytes (Figure 2C and 2E). Those long-term cultured cerebral organoids represent an innovative tool to study the genetic influence of patients with MS on oligodendrocyte maturation and myelination capacity, and possibly better understand the phenotypic differences in MS.

In our previous work, we identified p21 as a protein of interest in the pathogenesis of progressive MS. Indeed, p21 is required for the differentiation of oligodendrocytes of its ability to control exit from the cell cycle, and p21 knockdown mice have hypomyelinated brain (Zezula et al., 2001). Additionally, p21 is an autoimmunity suppressor (Santiago-Raber et al., 2001) as p21-/- mice develop a lethal autoimmune syndrome characterized by the production of pathogenic autoantibodies (Santiuste et al., 2010).

However, little is known about the implication of other members of the CDK inhibitors on MS. p21 belongs to the Cip/Kip-family which also includes p27 and p57 (Sherr and Roberts, 1999). The other main CDK inhibitor family, the INKs family, includes p16, p15, p18 and p19 (Safwan-Zaiter et al., 2022). We decided to analyze p16, p57 and p27 expressions in our c-organoid model of MS. p16 (p16INK4A), encodes an inhibitor of cyclin-dependent kinases 4 and 6 and plays an important role in cellular aging and in premature senescence (Ressler et al., 2006). Senescence can be induced by several factors such as DNA damage, oxidative stress or neuroinflammation and has been associated with several neurodegenerative disorders including Alzheimer’s disease, Down syndrome and Parkinson’s disease (Martinez-Cue and Rueda, 2020; Safwan-Zaiter et al., 2022). Interestingly p16 is involved in neural progenitors senescence in PPMS brain (Nicaise et al., 2019), while no difference were detected in PBMCs in patient with MS compared to controls (Yang et al., 2024). p57 can bind and inhibit all cyclin/CDK complexes, with a lower affinity for cyclin B/CDK1 and cyclin D2/CDK6 complexes (Matsuoka et al., 1996). It plays a crucial role in maintaining the quiescence of neural stem cells. While the ablation of p57 initially stimulates stem cell proliferation and neurogenesis, it eventually leads to the exhaustion of the neural stem cell population (Furutachi et al., 2013). More importantly, p57 has been associated with MS as it can inhibit the oligodendrocyte differentiation in EAE mice (Kremer et al., 2009). In neural stem cells, p57 expression influences their fate by promoting differentiation into astrocytes, thereby reducing the oligodendrocyte lineage (Jadasz et al., 2012). p27 is abundant in quiescent (G0) and early G1 phases and binds to and inhibits cyclinE/CDK2. It has other essential functions such as proliferation, differentiation, and cell death (Abbastabar et al., 2018). Interestingly, p27 is required for the differentiation of oligodendrocyte precursors into myelinating oligodendrocytes through cell cycle withdrawal (Casaccia-Bonnefil et al., 1997; Larocque et al., 2005) and is also required to promote timely oligodendrogenesis during neonatal brain development (Jablonska et al., 2012). Furthermore, p27 is a positive regulator of Schwann cell differentiation in vitro (Li et al., 2011). It is also important to note that p27 has distinct functions depending on its subcellular localization. It has been described as a tumor suppressor when expressed in the nucleus and as an oncoprotein when expressed in the cytoplasm (Blagosklonny, 2002).

In our work we have not detected any difference in p16, p27 or p57 expression in MS organoids compared to control. Marker location, expression and cell morphology were nearly identical in every condition (Figure 3). Only p21 expression was significantly lower in MS organoids as previously described (Daviaud et al., 2023). However, when we performed the same analysis in IPSC derived NPCs, we detected an important change in p16 expression in PPMS organoids compared to control, similar to the result obtained by Nicaise’s team (Nicaise et al., 2019), while no difference was observed for p57, p27 or p21. This result seems to indicate that NPCs in vitro and NPCs growing in organoids have different regulations and behaviors that will need further characterization.

In our organoid model of MS, we observed that p21 is the only dysregulated CDK inhibitor in the Cip/Kip and INK families. Therefore, we decided to study p21 regulators E2F1 and PAK. The E2F family of transcription factors plays a critical role in cell proliferation, differentiation, and apoptosis through transcription regulation (Muller et al., 2001). It is important to note that E2F1 is required for the activation of the p21 gene through Ras (Gartel et al., 2000; Radhakrishnan et al., 2004) and can act as a stemness regulator in neural stem cells, controlling neuronal cell numbers in the brain (Cooper-Kuhn et al., 2002). Interestingly, E2F1 also regulates the transition of oligodendrocyte precursors from proliferation to differentiation into oligodendrocytes, thereby stimulating myelination (Magri et al., 2014). Moreover, a link between E2F1 and MS has been described. Natalizumab, a FDA-approved treatment for MS induces the downregulation of miR-17, leading to an upregulation of E2F1 and p21 target genes (Meira et al., 2014). We did not observe any difference in E2F1 expression at d42, but did see a significant decrease in E2F1 expression at d120 in MS organoids, particularly PPMS, compared to control (Figure 5). These results suggest that E2F1 may not be involved in stem cell dysregulation, or cell cycle exit, but may play a role in later stages, such as oligodendrocyte differentiation and maturation in our MS paradigm. The dysregulation of oligodendrocyte differentiation and maturation may be a synergy between p21 and E2F1 dysregulation, as both proteins are essential for proper oligodendrocyte maturation and myelination, but also because E2F1 can control p21 expression.

p21-activated kinases (PAKs) are Cdc42/Rac–activated serine-threonine protein kinases that regulate several key pathways and are a major driver of neuronal development in humans. PAK1 is also implicated in several neurodevelopmental and neurodegenerative diseases, including autism, intellectual disability, and Alzheimer’s disease (Pan et al., 2015). PAK1 can also positively regulate oligodendrocyte differentiation and is required for the correct formation of myelin sheaths in the CNS (Brown et al., 2021). Moreover, it has been shown that PAK1 can block p21 expression (Prudnikova et al., 2016). We detected a significant increase of PAK1 expression at d42, particularly in PPMS and RRMS compared to control. Together those results suggest that while PAK1 increase could lead to an increase of oligodendrocyte differentiation, the PAK1 induced blockage of p21 expression may inhibit OPCs differentiation in oligodendrocyte, associated with the decrease of E2F1 expression which is also necessary for proper oligodendrocyte differentiation.

Previously, we showed a decrease of OPCs marker Olig2 in MS organoids compared to control cultured for 42 days (Daviaud et al., 2023). In our current work we decided to culture organoids up to 120-150 days in vitro to investigate mature and myelinating oligodendrocytes. In control organoids, we observed colocalization of the myelin marker MBP and the neuronal marker DCX indicating mature myelination of neurons. Interestingly, we detected a decrease in mature and myelinating oligodendrocytes in MS organoids, especially in PPMS (Figure 6). In parallel, we detected a significant decrease in p21 expression at d120 in PPMS, similar to our observations at d28 and d42. This work suggests that the defect in the cell cycle and of oligodendrocyte differentiation observed at d42 (Daviaud et al., 2023) leads to a significant decrease in oligodendrocyte maturation and myelination capacity which may explain the failure of remyelination observed in PPMS.

At d120, this organoid model of MS allowed us to study other neural cells that were not mature at d42 such as neurons and astrocytes. Astrocytes are active players in the pathogenesis of MS. Astrocytic dysfunction, associated with a MS genetic risk variant, suggests that astrocytes are involved in providing peripheral immune cells access to the central nervous system and that astrocyte-mediated processes are causative in lesion pathology (Ponath et al., 2018). In our research, we detected a significant decrease in astrocyte marker intensity in MS organoids, suggesting a decrease in glial cell differentiation (Figure 7). However, it has been suggested that reactive astrocytes change their morphology and size according to their reactive state in vitro and in vivo (Xiong et al., 2022), and thus the difference we detected in GFAP marker intensity could be due to a decrease in astrocyte size after activation in a pro-inflammatory mode. Therefore, further analysis of astrocyte properties in organoid model of MS is needed.

Our innovative model of MS using cerebral organoids at d120 also allowed the analysis of mature neurons such as GABAergic and glutamatergic neurons. While we did not observe any important change in GABAergic neuron population, we detected a significantly higher expression of glutamatergic neuron markers in PPMS and RRMS compared to control (Figure 7). Dysfunction of the GABAergic/glutamatergic pathways has already been reported in MS. Abnormal levels of glutamate have been reported in MS patients with increased levels in active lesion (Srinivasan et al., 2005) while a reduced concentration of GABA has been associated with cognitive impairment, physical disability, and fatigue in patients with MS (Cawley et al., 2015; Cao et al., 2018; Arm et al., 2021). Interestingly, astrocytes can regulate neurotransmitter homeostasis, as they uptake released neurotransmitters, such as glutamate and GABA, and then protects neurons from excitotoxicity and cell death (Mahmoud et al., 2019). In our organoid model, we detected a decrease in astrocyte population associated with an increase in glutamatergic neuron population. These results may suggest an important underlying mechanism for the clinical manifestations of cognitive impairment, physical disability or fatigue described in patients with MS.

Spinal cord abnormalities are common in MS and include a variety of pathological processes, such as demyelination, neuroaxonal loss and gliosis. Ultimately these result in motor weakness associated with movement difficulties, sensory disturbances, as well as bladder and bowel dysfunction (Walker, 2000). Spinal cord is affected by both inflammatory and neurodegenerative changes in patients with MS. We created spinal cord organoids from healthy control as well as patients with MS to have a more comprehensive review of MS pathogenesis and development. We observed a similar phenotype in our SCO and our c-organoids from patients with MS, with an increase of neurogenesis which may lead to an imbalance of excitatory and inhibitory neurons. We also checked for spinal cord motor neurons as motor neuron were reduced in postmortem MS patients (Vogt et al., 2009) but also in EAE rodent model (Gilmore et al., 2009) which may contribute to MS pathology. However, we didn’t detect any change in ChAT^+^ motor neurons in our SCO, meaning that the motor neuron degeneration may be inflammatory. Finally, we observed a decrease in astrocyte and oligodendrocyte populations, particularly in PPMS organoids, associated with a decrease in myelinating oligodendrocytes. Interestingly, a significant decrease in p21 expression was also detected in PPMS and RRMS organoids, demonstrating that the brain and spinal cord may share a similar pathophysiology in MS.

Taken together those results show that cerebral and spinal cord organoid are useful tools to study MS. At early time points it can be used to study precursor cell proliferation and differentiation capacity, while at later time points it can be used to study neural and glial cell maturation and even myelination capacity. We were able to confirm the involvement of the p21 pathway in MS and its associated regulators such as E2F1 and PAK1. The dysregulation of this protein expression leads to a significant decrease in oligodendrocyte differentiation, maturation, and myelination capacity, but also induces a defect of astrocyte population and seems to trigger an imbalance of inhibitory/excitatory neurons, which are all key factors in MS onset and symptom pathogenesis.

## ACKNOWLEDGMENTS.

The authors thank the members of Tisch MSRCNY for helpful discussions, and the New York Stem Cell Foundation for providing patients with MS derived IPS cell lines.

## Author contributions: CRediT taxonomy

Conceptualization, N.D., S.A.S. ; Methodology, N.D. ; Validation, N.D. ; Formal Analysis, N.D., T.M., W.H., A.M. ; Investigation, N.D., T.M., W.H., A.M. ; Writing – Original Draft, N.D. ; Writing – Review & Editing, N.D., S.A.S. ; Supervision, S.A.S. ; Project Administration, N.D. ; Funding Acquisition, N.D., S.A.S.

